# Predictive Power of Machine Learning Models for Relapse Outcomes in Acute Myeloid Leukemia: Unveiling Key Genes and Pathways for Improved Patient Management

**DOI:** 10.1101/2023.12.05.570249

**Authors:** Mehran Radak, Hossein Fallahi, Keyvan Karami

## Abstract

Acute Myeloid Leukemia (AML) is a challenging form of blood cancer requiring accurate relapse prediction for effective therapy and patient management. In this study, we applied multiple machine learning techniques to a dataset of AML patients in order to develop a reliable model for predicting relapse and guiding treatment decisions. We utilized various feature selection methods to identify the most relevant features associated with relapse. Additionally, we investigated gene ontology using the Gene Ontology (GO) database to gain insights into the biological processes and KEGG pathways related to the selected features. Our findings revealed key genes and pathways implicated in AML relapse. Among the machine learning models, Decision Tree (DT) showed the highest accuracy in predicting relapse outcomes. Furthermore, we compared the performance of DT models across different feature selections, highlighting the significance of specific factors such as MCL1, WBC, HGB, and BAD.p112 in relapse prediction. The results of our study have important implications for tailoring treatment plans and improving patient outcomes in AML. By accurately identifying patients at high risk of relapse, our model can aid in early interventions and personalized therapies. Ultimately, our research contributes to advancing the field of machine learning in AML and lays the foundation for developing effective strategies to combat relapse in this disease.

## Introduction

Acute myeloid leukemia (AML) is a type of blood cancer that affects the bone marrow and blood cells [1]. It is characterized by abnormal cell growth that interferes with producing healthy blood cells [2]. AML is the most common type of acute leukemia in adults, and its incidence increases with age. AML can develop rapidly and requires prompt diagnosis and treatment [3]. Symptoms may include fatigue, weakness, fever, easy bruising or bleeding, and frequent infections [4]. Diagnosis is typically made through a combination of blood tests, bone marrow biopsy, and imaging studies [5]. The treatment of AML usually involves chemotherapy, which aims to kill cancer cells and restore healthy blood cell production. In some cases, a bone marrow transplant may be necessary to replace the diseased bone marrow with healthy donor cells. Treatment choice depends on several factors, including the patient’s age, overall health, and the specific genetic characteristics of the cancer cells [6].

Despite advances in treatment, AML remains a challenging disease to manage, particularly in older adults. Relapse rates are high, and many patients require multiple rounds of treatment. As a result, there is a need for new and more effective therapies to improve outcomes for patients with AML [7]. Relapse occurs when cancer cells that were not eliminated by treatment start to grow and divide again. Several factors, including age, overall health, and specific genetic characteristics of the cancer cells, influence the risk of relapse [8]. AML is often treated with chemotherapy, which aims to eliminate cancer cells and restore healthy blood cell production. However, some cancer cells may survive the initial treatment and remain dormant in the body, which can lead to relapse [9]. In addition, the genetic mutations that drive the growth of cancer cells can change over time, making the cancer cells resistant to previously effective treatments [10]. The risk of relapse after initial treatment for AML is high, particularly in older adults. According to some estimates, up to 70% of patients with AML who achieve remission after initial treatment will eventually experience relapse [8]. To reduce the risk of relapse, some patients may be offered additional treatment after achieving remission. This may involve more intensive chemotherapy or a stem cell transplant to replace the diseased bone marrow with healthy donor cells. However, these treatments can be associated with significant side effects and may not be suitable for all patients [11].

Machine learning (ML) techniques have emerged as a valuable tool for predicting disease outcomes, including in the field of oncology [12]. ML algorithms use statistical models to learn patterns in large datasets and can be used to analyze a wide range of data types, including genomic, clinical, and imaging data [13]. In oncology, ML algorithms are being used to predict disease outcomes in various types of cancer, including acute myeloid leukemia (AML) [14]. ML algorithms can be used to analyze genomic data to identify specific genetic mutations associated with a higher risk of relapse in AML [15]. One of the main advantages of ML algorithms is their ability to analyze complex data sets and identify patterns that traditional statistical methods may miss. ML algorithms can identify subtle relationships between variables that may not be immediately apparent to human analysts. This can lead to more accurate predictions of disease outcomes and more personalized treatment plans for individual patients [16, 17]. As the field of ML advances, these algorithms will likely become increasingly important tools for predicting disease outcomes in AML and other types of cancer. The use of machine learning algorithms in predicting disease outcomes is a rapidly growing area of research. Researchers have used a variety of machine learning techniques, including artificial neural networks (ANN), decision trees (DT), logistic regression (LR), support vector machine (SVM), naive bayes (NB), random forest (RF), and multilayer perceptron (MLP) to predict cancer survival [18].

Interestingly, Random Forest (RF) appears to be the most preferred algorithm in most clinical studies. This is likely due to its ability to handle large and complex datasets while being less prone to overfitting than other machine learning algorithms. However, it is important to note that the choice of an algorithm may depend on the specific characteristics of the dataset and the research question being investigated [19].

The current study evaluates the predictive power of multiple machine learning techniques applied to a dataset of patients with Acute Myeloid Leukemia (AML) in predicting relapse outcomes. We aim to improve the subsequent therapy and management of AML patients by identifying those at high risk of relapse and intervening early to decrease the risk. We utilized various machine learning models to assess their accuracy in predicting the relapse of AML patients based on clinical data. Our study builds on previous research that has shown the effectiveness of machine learning algorithms in predicting survival in this cancer [14]. Our ultimate goal is to develop a reliable and accurate model that can be used in clinical practice to guide treatment decisions and improve patient outcomes. By accurately predicting the risk of relapse, we can tailor treatment plans to each patient’s specific needs and intervene early to prevent or manage relapse.

## Methods

In this study, we utilized a database containing various clinical variables of patients with AML, including numerical and categorical data. The patients were classified according to the French-American-British (FAB) system. Data preparation for analysis was performed using data mining tools and algorithms. We applied feature selection via feature weighting methods to identify the most important features in predicting relapse outcomes. A total of 25 high-weight features were selected to continue the analysis. Multiple classifiers were then trained and evaluated to determine their ability to predict relapse outcomes in AML patients accurately. The performance of each classifier was evaluated using various metrics such as accuracy, sensitivity, specificity, and area under the receiver operating characteristic curve (AUC-ROC). To ensure the validity of our results, we applied a ten-fold cross-validation technique and repeated the analysis on multiple subsets of the data. Our methodology was designed to identify the most accurate and robust classifier for predicting relapse outcomes in AML patients based on clinical data.

### Dataset

The data utilized in this study were obtained from the Leukemia Sample Bank at the University of Texas M. D. Anderson Cancer Center. The primary dataset consisted of information collected between January 15, 1998, and March 9, 2006, pertaining to 249 patients diagnosed with AML [20]. Multiple variables were evaluated for each patient, including both categorical and numerical data. To ensure that the data were suitable for analysis, missing values in categorical variables and numerical features were replaced using the mode and average of the missing value in each class.

This process was performed using the RapidMiner software (version 10.1.002, www.rapidminer.com).

### Data cleaning

The first step in our data analysis was to preprocess the data to ensure it was suitable for analysis. This process, commonly known as data cleaning, involves removing any irrelevant or correlated attributes that could negatively impact the model’s accuracy. To this end, we removed correlated attributes with a Pearson correlation coefficient greater than 0.95 from the dataset. Additionally, we identified numerical attributes with a standard deviation less than or equal to a given threshold of 0.1 and removed them from the initial dataset as they were deemed useless in the analysis. The remaining data, which had undergone this preprocessing, was then treated as the processed dataset and used for our study. By removing any irrelevant or correlated attributes, we increased the model’s predictive ability and obtained a better fit of the model.

### Feature selection

Feature selection is an important step in machine learning to obtain a good model fit and improve predictive ability. This study used feature weighting approaches, including information gain, information gain ratio, Gini index, chi-squared, correlation, relief, and uncertainty, to select relevant features. We selected 25 features with the highest weighting score for further analysis, representing more than 30% of all features analyzed.

Here is a brief description of the feature selection algorithms used in this study:

1. Information gain: An entropy-based feature evaluation method that measures the amount of information the feature items provide for the text category. It is widely used in machine learning for feature selection [21].
2. Information gain ratio: The information gain ratio is the gain divided by the entropy of A for splitting according to some feature “A” [22].
3. Gini index: A split measure of total variance across the K classes used for splitting attributes in decision trees [23].
4. Chi-squared: A statistical technique used to test the independence of two events. In feature selection, it is used to test the independence of the term’s occurrence and the class’s occurrence [24].
5. Correlation: A statistical method used to measure the linear relationship between two variables. Highly correlated features have almost the same effect on the dependent variable, so when two features have a high correlation, one of them can be dropped [25].
6. Relief: A successful feature selection algorithm that estimates the quality of features based on how well their values distinguish between the instances of the same and different classes. It was originally defined for two-class problems and later extended to handle noise and multiclass datasets [26].
7. Uncertainty: Symmetric uncertainty measures the correlation between two variables by normalizing mutual information to the entropies of two variables. In feature selection, it evaluates an attribute by measuring its symmetrical uncertainty concerning the class [27].

### Gene Ontology Analysis

A key aspect of our study involved performing Gene Ontology (GO) analysis to gain insights into the functional properties of genes and gene products across different species. The aim was to standardize the representation of these properties and understand their relevance to our investigation of Acute Myeloid Leukemia (AML) relapse outcomes. To conduct the GO analysis, we utilized the online DAVID (Database for Annotation, Visualization, and Integrated Discovery) tool.

Our focus was specifically on investigating biological processes (BP) and pathways from the Kyoto Encyclopedia of Genes and Genomes (KEGG). In this analysis, we selected the terms that had the highest number of associated genes and a p-value below 0.05. This ensured that we included the most significant and relevant terms for our study. By identifying these enriched terms, we aimed to uncover the biological processes and pathways that were closely linked to the prediction of relapse outcomes in AML. The GO analysis provided valuable insights into the functional roles and relationships of genes and their involvement in key biological processes and pathways. These findings further supported our understanding of the underlying mechanisms associated with AML relapse and contributed to the overall interpretation and discussion of our research results.

### Model evaluation

This study evaluated the performance of six machine learning techniques (RF, DT, LR, Naive Bayes, W-Bayes Net, and GBT) using 10-fold cross-validation on multiple datasets. The conventional machine learning algorithms were assessed based on their accuracy, kappa, sensitivity, specificity, positive predictive value, negative predictive value, and AUC. The ROC curve was used to plot the true positive rate (TPR) against the false positive rate (FPR) at various threshold settings, and the AUC was calculated as the area under the ROC curve. A model with an AUC closer to 1 predicted outcomes better. The abbreviations used for empirical quantities in the measurements were P (# positive samples), N (# negative samples), TP (# true positives), TN (# true negatives), FP (# false positives), and FN (# false negatives). Accuracy was estimated using the ratio of (TP+TN)/(P+N), while positive predictive value (PPV) was estimated by TP/(TP+FP) and negative predictive value (NPV) by TN/(TN+FN). Sensitivity was estimated by TP/P and specificity by TN/N. Accuracy was used to select the optimal model with the highest value. Sensitivity in this context is also referred to as the true positive rate or recall, and PPV is also referred to as precision.

The six machine learning techniques used in this study were Random Forest (RF), Decision Tree (DT), Logistic Regression (LR), Naive Bayes, Bayesian Network (Bayes Net), and Gradient Boosted Tree (GBT). RF creates multiple classification and regression (CART) trees, each trained on a bootstrap sample of the original training data, and searches across a randomly selected subset of input variables to determine the split [28]. DT constructs decision trees that are relatively fast to build and produce interpretable models [29]. LR is a technique borrowed from statistics, suitable for regression analysis with a dichotomous dependent variable [30]. Naive Bayes is a predictive model and classification technique based on the Bayes theorem, assuming that all predictors are independent [31]. Bayes Net is a probabilistic graphical structure representing the probabilistic relationships between variables via a directed acyclic graph. GBT trains a model by iteratively improving a single tree model, with the final model being a weighted sum of all created models [32].

## Results

### Feature selection

Various feature selection operators were used to select the top 25 features, which accounted for approximately 30% of all features. These features, along with the most important feature that had a higher weight score in multiple feature selection algorithms, are presented in Table 1.

**Table 1.**
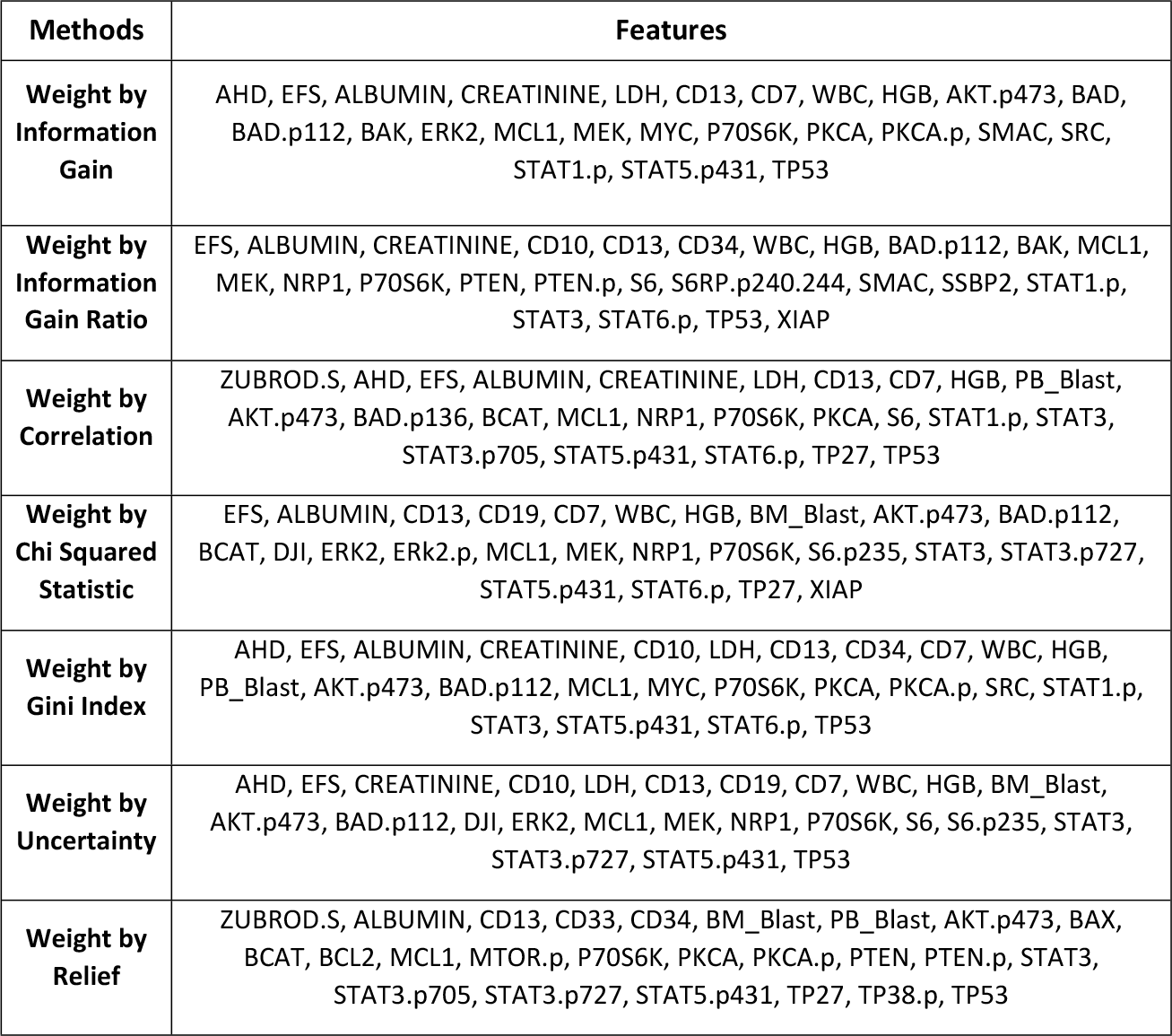
The top 25 selected features using Different Feature Selection Methods.

Among the seven feature selection techniques employed in our study, we identified several common features that consistently showed high relevance in predicting relapse outcomes in Acute Myeloid Leukemia (AML) patients. These features, as depicted in Table 2, include CD13, MCL1, P70S6K, HGB, EFS, ALBUMIN, TP53 and etc. These features were selected using various techniques such as Information Gain, Information Gain Ratio, Gini Index, Chi-Squared, Correlation, and Relief. The presence of these features in the selected set highlights their potential significance in contributing to the relapse of AML. CD13, MCL1, and TP53 have been widely studied in the context of AML and are known to play crucial roles in disease progression and treatment response. P70S6K, HGB, and ALBUMIN are indicators of cellular processes, red blood cell count, and nutritional status, respectively, which have been associated with disease outcomes in AML. EFS (Event-Free Survival) represents the time period without disease recurrence or other defined events, while WBC (White Blood Cell count) and BAD.p112 (a protein associated with apoptosis regulation) reflect aspects of the immune system and cell signaling pathways, respectively. The inclusion of these features in our predictive models underscores their potential as valuable predictors for relapse risk in AML. By incorporating these factors into our machine learning models, we improve the accuracy and reliability of our predictions, enabling clinicians to identify patients at high risk of relapse and intervene early to optimize treatment strategies. The identification of these common features emphasizes the importance of a comprehensive and multi-dimensional approach to feature selection in AML prediction models. By considering a wide range of clinical and molecular factors, we can better understand the underlying mechanisms of relapse and develop more robust and accurate models for personalized treatment decision-making in AML patients.

**Table 2.**
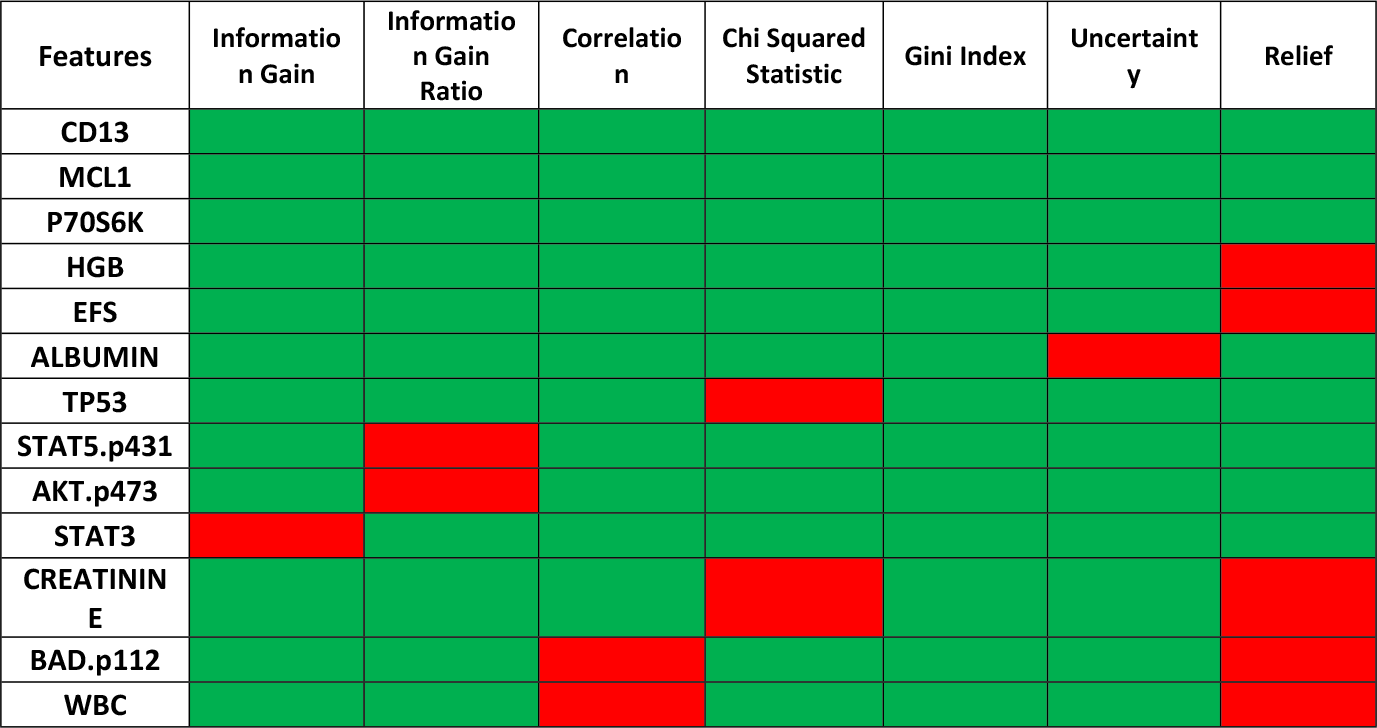
Most common features Associated with Each Feature Selection Method. Green square: indicates the existence of the feature. Red square: indicates the absence of the feature.

### Gene ontology for each feature

The investigation of gene ontology was carried out using the David database, allowing us to explore the functional relevance of the features identified in our study. By analyzing the number of genes associated with each feature, we gained insights into the molecular mechanisms underlying relapse in Acute Myeloid Leukemia (AML). The details of this investigation are presented in Table 3, which provides information on the genes associated with each feature using various feature selection methods.

**Table 3.**
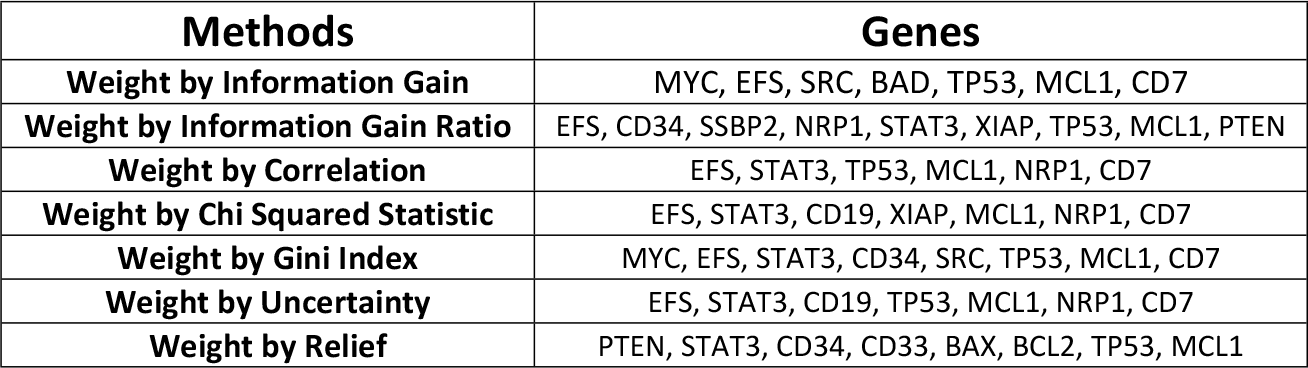
Genes Associated with Each Feature Using Different Feature Selection Methods.

It is worth noting that the identified genes may have implications for disease prognosis, therapeutic response, and potential targets for intervention. However, further experimental validation and functional characterization of these genes are warranted to establish their precise roles in AML relapse and determine their potential clinical significance. By integrating machine learning techniques with gene ontology analysis, our study provides a comprehensive understanding of the predictive power of different features and their associated genes in predicting relapse outcomes in AML. These findings contribute to the development of accurate predictive models and aid in the identification of high-risk patients who may benefit from early intervention strategies, ultimately leading to improved management and better patient outcomes.

Based on the analysis of the genes obtained through each feature selection method, we identified the biological processes and KEGG pathways associated with each method. The results are summarized in Table 4.

**Table 4.** Biological Processes and KEGG Pathways associated with each Feature Selection Method.

| **Methods** | Most number of genes involved in **BP** | **BP pathways** | Most number of genes involved in **KEGG** | **KEGG pathways** |
| --- | --- | --- | --- | --- |
| **Information Gain** | 4 | cellular response to hypoxia/ response to xenobiotic stimulus/ positive regulation of apoptotic process/ negative regulation of apoptotic process | 4 | Thyroid hormone signaling pathway/ Hepatitis B/ PI3K-Akt signaling pathway |
| **Information Gain Ratio** | 4 | negative regulation of apoptotic process/ regulation of cell cycle | 4 | Human T-cell leukemia virus 1 infection/ MicroRNAs in cancer/ Pathways in cancer |
| **Correlation** | 2 | negative regulation of autophagy/ positive regulation of angiogenesis | 3 | MicroRNAs in cancer |
| **Chi Squared Statistic** | 2 | negative regulation of autophagy/ positive regulation of angiogenesis | ----- | ----- |
| **Gini Index** | 4 | positive regulation of gene expression/ negative regulation of apoptotic process | 4 | Kaposi sarcoma-associated herpesvirus infection/ Proteoglycans in cancer/ MicroRNAs in cancer |
| **Uncertainty** | 2 | negative regulation of autophagy/ positive regulation of angiogenesis | 3 | MicroRNAs in cancer/ PI3K-Akt signaling pathway |
| **Relief** | 5 | apoptotic process | 5 | MicroRNAs in cancer/ Pathways in cancer |

The chart (Figure 1) illustrates the number of most common biological processes (BP) and KEGG pathways associated with each feature selection method. This chart visually demonstrates the distribution of the most common pathways across different feature selection methods, providing insights into the functional relevance of the selected genes in AML relapse.

**Figure 1.**
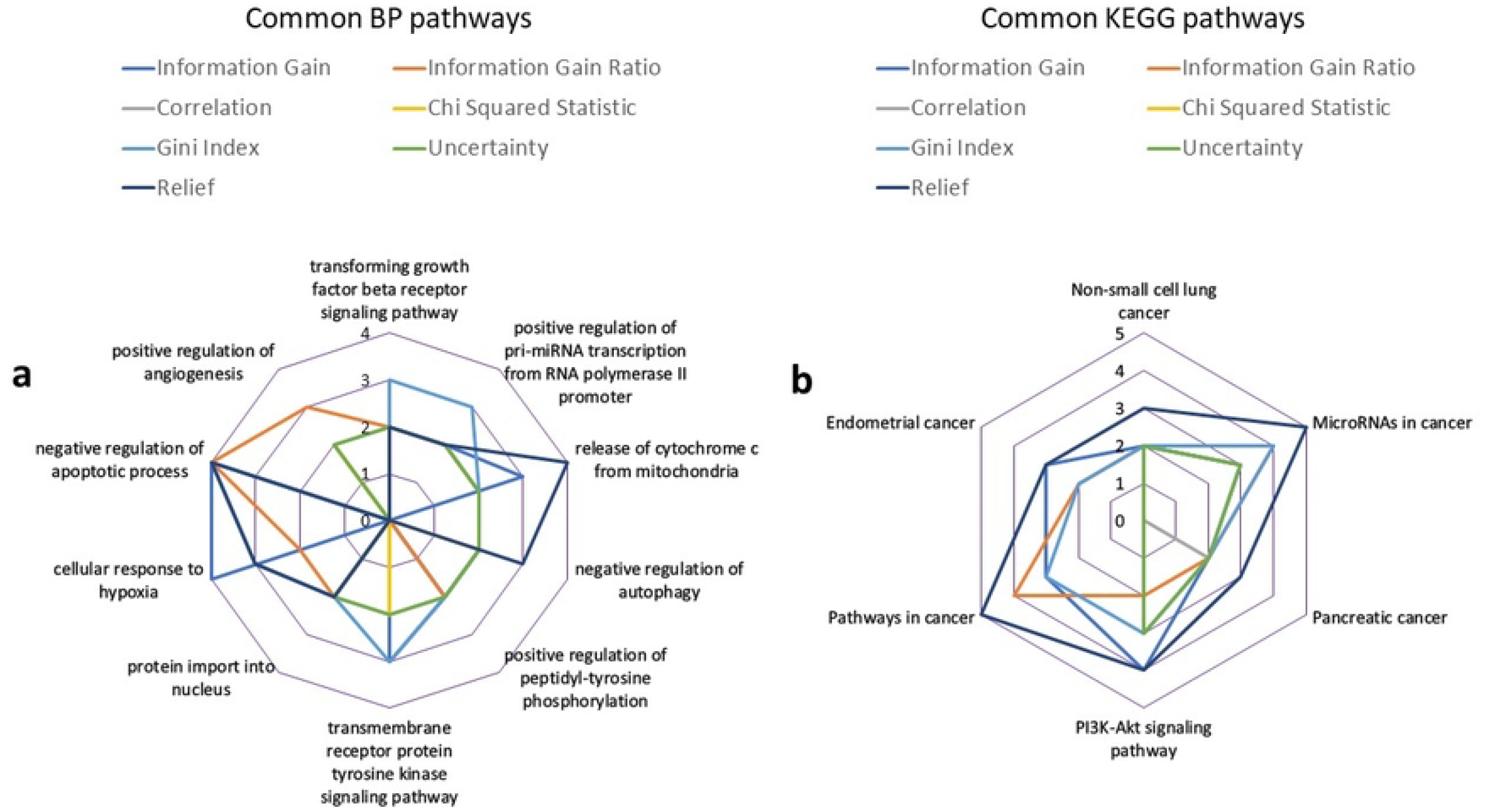
The number of Most Common Biological Processes (BP) and KEGG Pathways for each Feature Selection Method.

### Assessing the predictive ability of the model

Table 5 presents the accuracy of each machine learning model based on different feature selection methods. The models evaluated include Random Forest (RF), Decision Tree (DT), Logistic Regression (LR), Naive Bayes (NB), and Gradient Boosted Tree (GBT). For each feature selection method, the accuracy of the models is shown as a percentage. It can be observed that the performance of the models varies across the different feature selection methods. It is evident that the accuracy of the models varies across the different feature selection methods. This suggests that the choice of feature selection technique plays a significant role in determining the predictive performance of the models. The Chi Squared Statistic method consistently achieves high accuracy across most models, making it a promising approach for feature selection in this context. The RF and Gini index methods also perform well, demonstrating their effectiveness in capturing relevant information for prediction. On the other hand, the Correlation and Uncertainty methods exhibit relatively lower accuracies compared to other techniques. While they may still provide valuable insights, it may be worthwhile to explore alternative feature selection methods to potentially improve the models’ performance. These findings highlight the importance of careful feature selection in developing accurate predictive models. By selecting the most informative features, we can enhance the models’ ability to capture meaningful patterns and make reliable predictions regarding relapse outcomes in patients with Acute Myeloid Leukemia (AML). The results suggest that the choice of feature selection method can impact the accuracy of the machine learning models. Further analysis and comparison of these models can help identify the most effective approach for predicting relapse outcomes in patients with Acute Myeloid Leukemia (AML).

**Table 5.**
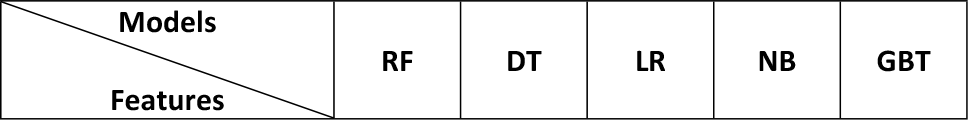
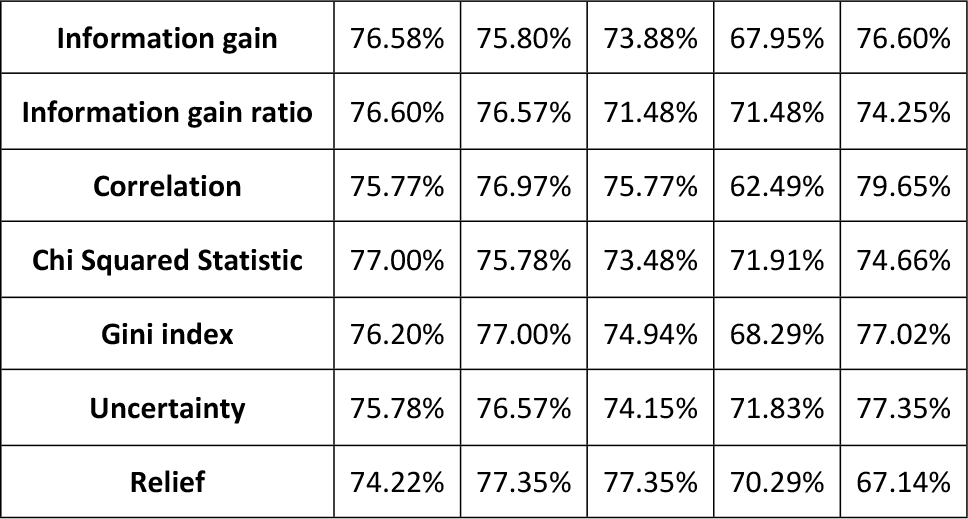
Accuracy of Each Model Based on Feature Selection Methods.

| Models<br>Features | RF | DT | LR | NB | GBT |
| --- | --- | --- | --- | --- | --- |
| **Information gain** | 76.58% | 75.80% | 73.88% | 67.95% | 76.60% |
| **Information gain ratio** | 76.60% | 76.57% | 71.48% | 71.48% | 74.25% |
| **Correlation** | 75.77% | 76.97% | 75.77% | 62.49% | 79.65% |
| **Chi Squared Statistic** | 77.00% | 75.78% | 73.48% | 71.91% | 74.66% |
| **Gini index** | 76.20% | 77.00% | 74.94% | 68.29% | 77.02% |
| **Uncertainty** | 75.78% | 76.57% | 74.15% | 71.83% | 77.35% |
| **Relief** | 74.22% | 77.35% | 77.35% | 70.29% | 67.14% |

### Choosing best model

In order to select the best model for predicting relapse in Acute Myeloid Leukemia (AML) patients, you employed the Receiver Operating Characteristic (ROC) curve analysis. The ROC curve is a widely used tool for evaluating the performance of classification models. Based on your analysis, the Decision Tree model emerged as the best-performing model among the ones evaluated. The ROC curve comparison revealed that the Decision Tree model achieved the highest Area Under the Curve (AUC) value, indicating its superior discriminatory power in distinguishing between relapse and non-relapse cases. The high AUC value suggests that the Decision Tree model has a strong ability to correctly classify patients with AML who are at high risk of relapse. This finding demonstrates the model’s potential for accurately identifying individuals who require closer monitoring and early intervention to mitigate the risk of relapse. The selection of the Decision Tree model as the best model highlights its capability to capture relevant patterns and make accurate predictions based on the clinical data available. Its inherent ability to create decision rules and hierarchically partition the feature space likely contributes to its strong predictive performance in this context. It is important to note that while the Decision Tree model has shown promising results in predicting relapse outcomes, further validation and evaluation are necessary. Robust statistical analysis and validation on independent datasets would enhance the reliability and generalizability of the findings.

Additionally, considering the complexity and heterogeneity of AML, a combination of multiple models or ensemble techniques could potentially improve predictive accuracy further.

### Decision tree models

Figure 3 illustrates the Decision Tree models generated using different feature selections. Each model represents a specific set of features that were identified as important for predicting relapse outcomes in Acute Myeloid Leukemia (AML) patients. The Decision Tree algorithm was applied to each feature selection to create a predictive model. Each model in Figure 3 visually represents the decision-making process of the Decision Tree algorithm, with nodes representing feature splits and branches leading to different predictions or outcomes. The structure and patterns of the Decision Trees provide insights into how each feature contributes to the prediction of relapse in AML patients. A comparative analysis of the Decision Tree models based on different feature selections allows researchers and clinicians to understand the distinct contributions of MCl1, WBC, HGB, and BAD.p112 in predicting relapse outcomes in AML.

**Figure 2.**
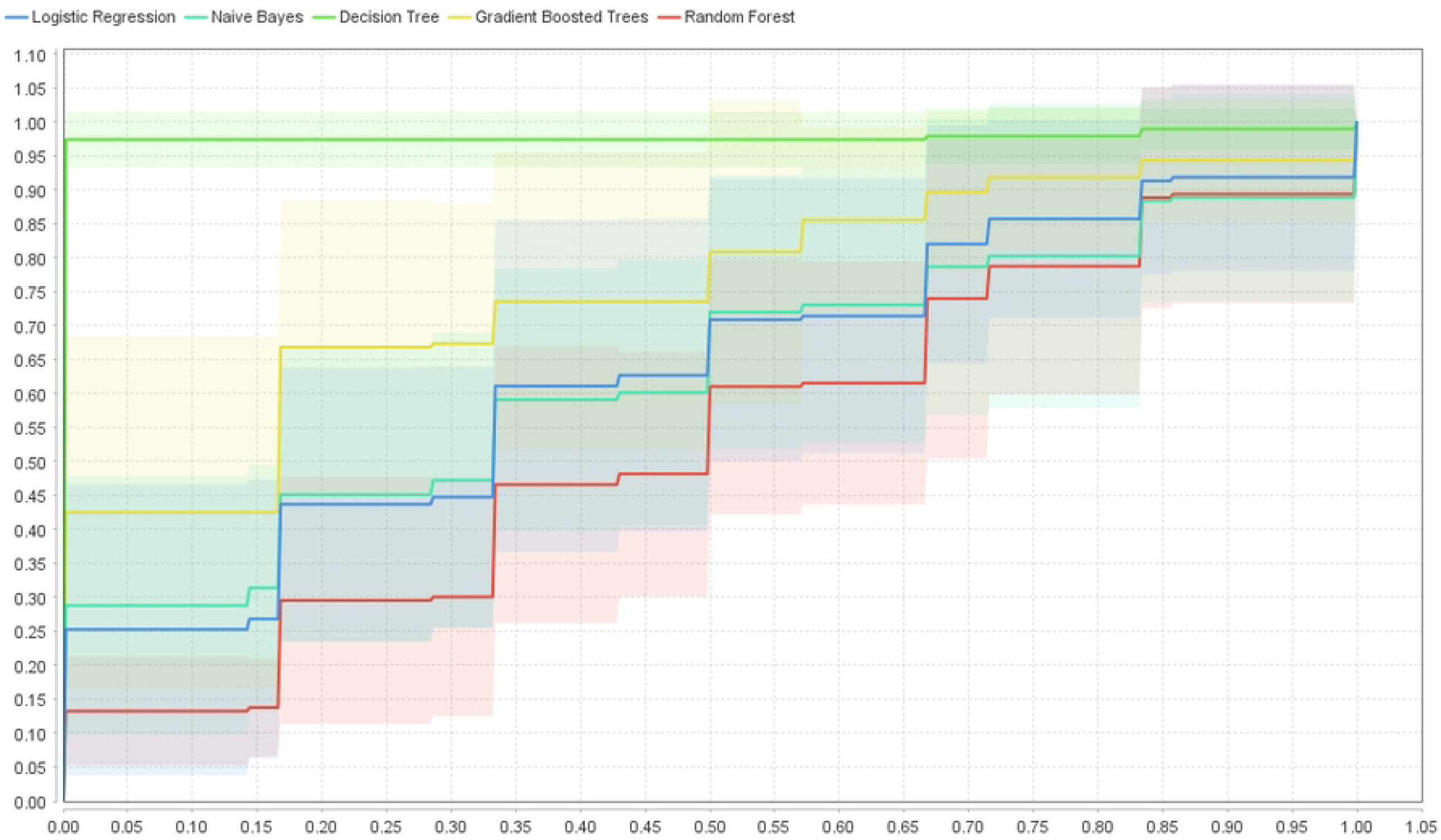
Represents the ROC curve for each model.

**Figure 3.**
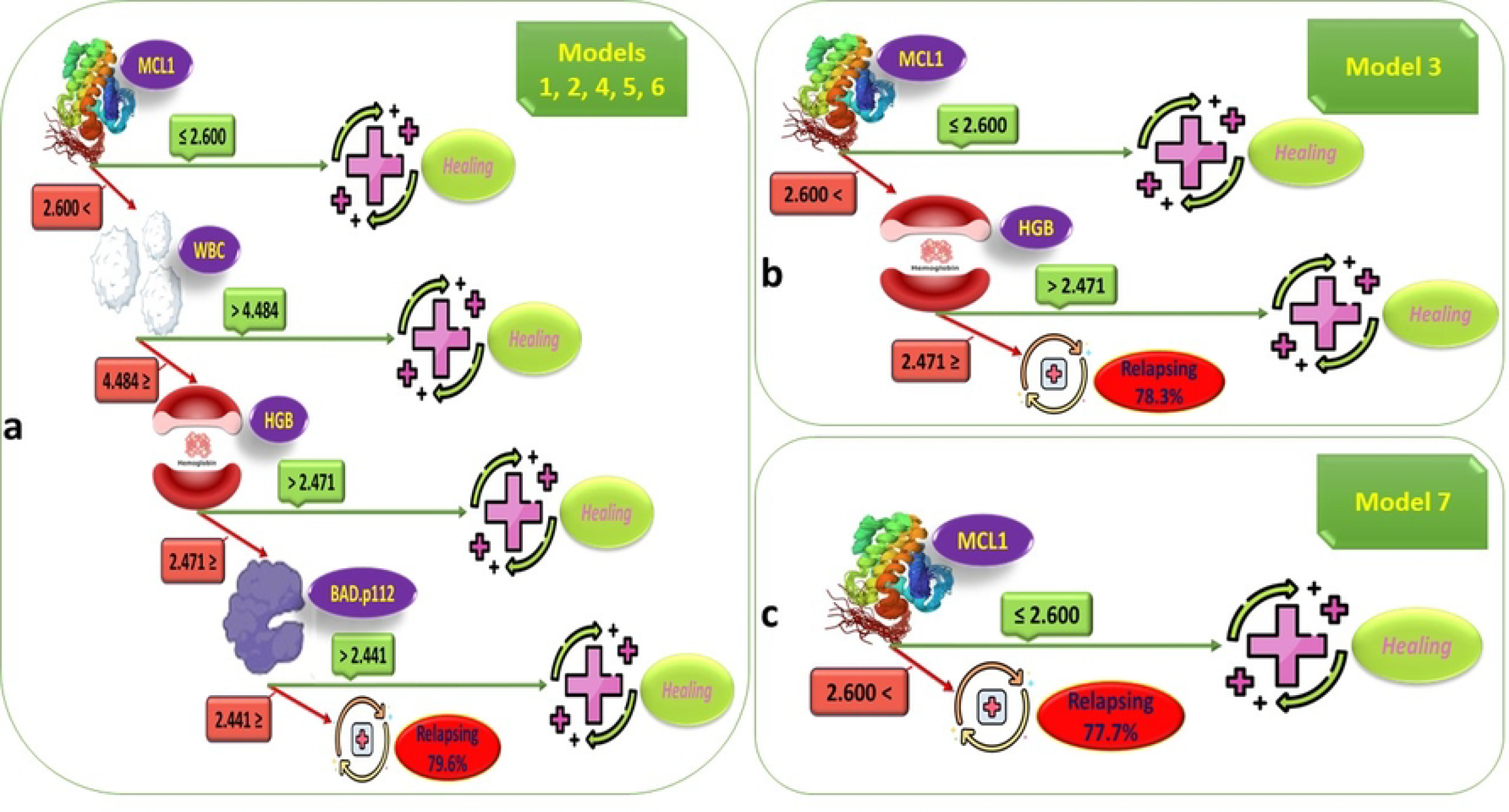
Decision Tree Models for Different Feature Selections.

## Discussion

Acute Myeloid Leukemia (AML) is a complex and aggressive form of blood cancer that requires comprehensive approaches for effective therapy and management. The identification of patients at high risk of relapse and the ability to intervene early are crucial factors in improving patient outcomes. In recent years, machine learning techniques have emerged as promising tools for predicting relapse outcomes and guiding treatment decisions in AML. In this study, we aimed to assess the predictive power of various machine learning models in determining the relapse outcomes of AML patients based on clinical data. Our research is built upon previous studies that have demonstrated the effectiveness of machine learning algorithms in predicting survival in AML [14]. We sought to develop a reliable and accurate model that could be utilized in clinical practice to tailor treatment plans and improve patient outcomes. To achieve our objectives, we employed a dataset of AML patients and applied multiple machines learning techniques, including Random Forest (RF), Decision Tree (DT), Logistic Regression (LR), Naive Bayes (NB), and Gradient Boosting Trees (GBT). We also employed feature selection methods, such as Information Gain, Information Gain Ratio, Correlation, Chi Squared Statistic, Gini Index, Uncertainty, and Relief, to identify the most relevant features associated with relapse. Our study included a comprehensive evaluation of each model’s accuracy in predicting relapse outcomes based on different feature selections. Furthermore, we explored the gene ontology using the Gene Ontology (GO) database and investigated the biological processes and KEGG pathways associated with the selected features. This analysis provided valuable insights into the functional roles and interactions of genes involved in AML relapse. In this study, we found a number of factors that we will discuss further in the text.

MCL1 (myeloid cell leukemia 1) is a protein that plays a crucial role in the regulation of cell survival and apoptosis. It has been identified as a factor contributing to the relapse of Acute Myeloid Leukemia (AML). MCL1 is part of the BCL2 family of proteins, which are involved in the regulation of cell death and survival. Overexpression or dysregulation of these proteins can lead to the development of cancer, including AML. In the context of AML, MCL1 has been found to be overexpressed in leukemic cells, contributing to their survival and resistance to chemotherapy. This overexpression can lead to a higher risk of relapse in AML patients, as the leukemic cells are more resistant to treatment. In a study examining the protein expression signatures in AML patients treated with venetoclax, a BCL2 inhibitor, alterations in MCL1 was noted at relapse, suggesting a role for MCL1 in the development of resistance to this targeted therapy [33]. One of the challenges in treating AML is the presence of leukemia stem cells (LSCs), which are believed to be the driving factor behind disease relapse and chemotherapeutic resistance. The development of 3D in vitro models for studying AML has shown promise in understanding the mechanisms of chemotherapeutic resistance in LSCs and identifying potential therapeutic targets, such as MCL1 [34]. MCL1 has been found to contribute to the relapse of the disease through various mechanisms. One way MCL1 contributes to AML relapse is by promoting resistance to certain treatments. For example, MCL1 has been found to be differentially expressed in triple-negative breast cancer (TNBC) tissues and cells, potentially limiting the efficacy of anticancer agents like recombinant human tumor necrosis factor-related apoptosis-inducing ligand (rh-TRAIL) [35]. In AML, MCL1 downregulation has been observed when CDK7 inhibitor THZ1 is combined with azacitidine (AZA), resulting in synergistic antileukemic effects in AML cell lines and primary cells [36]. This suggests that MCL1 plays a role in the resistance of AML cells to certain treatments, and its inhibition can enhance the effectiveness of these therapies. Additionally, MCL1 expression and function have been linked to the effectiveness of certain AML treatments. For instance, an alvocidib-containing regimen has been found to be highly effective in AML patients through a mechanism dependent on MCL1 expression and function. In summary, MCL1 is an anti-apoptotic protein that contributes to the relapse of AML by promoting resistance to certain treatments and playing a role in the effectiveness of specific therapies.

Targeting MCL1 and understanding its role in AML relapse could potentially lead to the development of more effective treatment strategies for patients with this aggressive form of leukemia.

WBC (White Blood Cell) count, a factor associated with relapse in Acute Myeloid Leukemia (AML), plays a crucial role in understanding disease progression and predicting patient outcomes. AML is a hematological malignancy characterized by the rapid proliferation of abnormal myeloid cells in the bone marrow, leading to an imbalance in blood cell production. In a study of pediatric patients with core binding factor-acute myeloid leukemia (CBF-AML), the initial WBC count was analyzed as a potential prognostic factor, although it was not identified as an independent risk factor affecting overall survival in this specific study [37]. However, in another study focusing on AML patients with t(8;21), the presence of KIT mutations at codon 816 was associated with a high WBC count at diagnosis, a high incidence of extramedullary leukemia, and a high risk of relapse during the course of the disease [38]. Moreover, a study on AML patients with deranged core-binding factor beta (CBFβ) found that increasing WBC count at diagnosis was an independent risk factor for relapse incidence, with a 1.1% increase in hazard for each 1.0 × 10^9/L WBC increment [39]. In another study, higher levels of phosphorylated FOXO3A, a tumor suppressor gene, were associated with increased proliferation, evidenced by a strong correlation with higher WBC count, percent marrow, and blood blasts [40]. The WBC count has been identified as a factor contributing to the relapse of AML in various studies, although its impact may vary depending on the specific subtype of AML and other factors, such as the presence of gene mutations.

Hemoglobin (HGB) is an essential protein found in red blood cells that carries oxygen from the lungs to the rest of the body. In the context of Acute Myeloid Leukemia (AML) relapse, HGB has been identified as a factor contributing to the relapse of the disease. In the context of AML, lower HGB levels have been associated with an increased risk of relapse. A decrease in HGB indicates a compromised red blood cell count and impaired oxygen-carrying capacity, which can have significant implications for disease progression and treatment outcomes. HGB levels may be an important factor to consider when assessing the risk of relapse and overall survival in AML patients. Regarding the role of HGB in other cancer relapses, there is limited information available. However, it is worth noting that HGB levels have been associated with the health status of patients undergoing treatment for various types of cancer, and low HGB levels have been linked to poor prognosis in some cases [41].

BAD.p112 is a member of the Bcl-2 protein family, which includes both pro-apoptotic and anti-apoptotic proteins. In cancer cells, dysregulation of the apoptosis pathway can contribute to treatment resistance and disease relapse. BAD.p112 is specifically involved in promoting apoptosis by binding to and inhibiting anti-apoptotic Bcl-2 family members, thereby allowing pro-apoptotic proteins to induce cell death [42]. This suggests that the activation of the apoptotic pathway through increased expression or activity of BAD.p112 may lead to enhanced cell death in leukemic cells, reducing the risk of relapse. Conversely, decreased expression or functional impairment of BAD.p112 can confer a survival advantage to leukemic cells, allowing them to evade apoptosis and persist in the bone marrow. This resistance to programmed cell death can contribute to disease relapse by enabling the survival and proliferation of residual leukemic cells following treatment. While the specific role of BAD.p112 in AML relapse is not well-documented, it is essential to continue researching and understanding the factors contributing to relapse in AML patients. By accurately predicting the risk of relapse, treatment plans can be tailored to each patient’s specific needs, and early interventions can be made to prevent or manage relapse, ultimately improving patient outcomes.

CD13 is a protein that has been found to be overexpressed in some cancer cells, including gastric cancer and has been shown to play a role in cancer metastasis. Inhibiting CD13 has been found to suppress the ability of migration and invasion in gastric cancer cells, suggesting that it may be a potential target for cancer therapy [43]. In hepatocellular carcinoma (HCC), CD13+ cancer stem cells have been found to drive relapse, and it has been suggested that targeting CD13 may be a potential therapeutic strategy [44]. TP53 is a tumor suppressor gene that plays a critical role in preventing the development of cancer. Mutations in TP53 are common in many types of cancer and are associated with poor prognosis and resistance to therapy. In prostate cancer, a transcriptomic signature for relapse prediction has been identified from the differentially expressed genes between TP53 mutant and wild-type tumors [45]. P70S6K is a protein that is involved in cell proliferation and survival and has been implicated in breast cancer recurrence. In breast cancer, interfering with p70S6K activity, which is downstream of STAT5, has been found to impair local relapse [46]. Cancer relapse can occur even decades after the primary therapy, and markers are needed to predict cancer progression and the risk of late recurrence. Low cytokeratin CK5 expression has been found to be associated with metastases in breast cancer, and CK5 positivity has distinguished triple-negative tumors into rapidly and slowly recurring cancers [47].

## Conclusion

In conclusion, our study focused on the predictive power of multiple machine learning techniques applied to a dataset of patients with Acute Myeloid Leukemia (AML) to improve the subsequent therapy and management of AML patients by identifying those at high risk of relapse. By accurately predicting the risk of relapse, our goal is to develop a reliable and accurate model that can be used in clinical practice to guide treatment decisions and improve patient outcomes. Through the utilization of various machine learning models and the analysis of clinical data, we have built upon previous research that has demonstrated the effectiveness of machine learning algorithms in predicting survival in AML. Our study contributes to the existing body of knowledge by focusing specifically on the prediction of relapse outcomes, a critical aspect of AML management.

We have identified several factors, including WBC, HGB, and BAD.p112, that significantly contribute to the relapse of AML. These factors play crucial roles in the disease progression and treatment response, highlighting their potential as valuable predictors of relapse risk. Integrating these factors into our predictive models has enhanced their accuracy in assessing the likelihood of relapse, enabling clinicians to tailor treatment plans to each patient’s specific needs and intervene early to prevent or manage relapse. Our findings underscore the importance of considering not only traditional clinical parameters but also molecular markers and signaling pathways involved in AML progression. Machine learning techniques provide a powerful tool to integrate and analyze these complex datasets, facilitating the development of reliable and accurate prediction models. The successful implementation of our predictive model in clinical practice could revolutionize the management of AML patients. By identifying those at high risk of relapse, clinicians can proactively intervene and optimize therapeutic strategies to decrease the risk and improve patient outcomes. Tailoring treatment plans based on individualized relapse risk assessments has the potential to enhance treatment efficacy, minimize unnecessary treatments, and reduce healthcare costs.

However, further research and validation are necessary to refine and validate our predictive model. Future studies should explore additional factors and refine the integration of clinical and molecular data to enhance the accuracy and generalizability of the model. Moreover, investigating the underlying mechanisms of the identified factors, such as the role of BAD.p112 in apoptosis regulation, may uncover new therapeutic targets for intervention and provide opportunities for personalized treatment approaches.

## Declarations

### Ethics approval and consent to participate

Not applicable

### Consent for publication

Not applicable

### Consent for parcipitaion

Not applicable

### Availability of data and material

We have obtained the original data from the following website: http://bioinformatics.mdanderson.org/Supplements/Kornblau-AML-RPPA/aml-rppa.xls.

### Code availability

The request for codes and models should be send to corresponding author.

### Authors’ contributions

MR and HF conceived the idea. MR and KK conducted the experiment and prepared the results. HF supervised the procedure and verified the results and discussion. MR wrote the first draft, and HF edited the manuscript. MR and HF prepared the final draft.

## Acknowledgments

Not applicable

## References

1. Pelcovits, A. and R. Niroula, Acute myeloid leukemia: a review. Rhode Island medical journal, 2020. 103(3): p. 38–40.

2. Kumar, C.C., Genetic abnormalities and challenges in the treatment of acute myeloid leukemia. Genes & cancer, 2011. 2(2): p. 95–107.

3. Shallis, R.M., et al., Epidemiology of acute myeloid leukemia: Recent progress and enduring challenges. Blood reviews, 2019. 36: p. 70–87.

4. Sultan, S., et al., Demographic and clinical characteristics of adult acute Myeloid Leukemia-tertiary care experience. Asian Pacific Journal of Cancer Prevention, 2016. 17(1): p. 357–360.

5. Percival, M.-E., et al., Bone marrow evaluation for diagnosis and monitoring of acute myeloid leukemia. Blood reviews, 2017. 31(4): p. 185–192.

6. Showel, M.M. and M. Levis, Advances in treating acute myeloid leukemia. F1000prime reports, 2014. 6.

7. DeWolf, S. and M.S. Tallman, How I treat relapsed or refractory AML. Blood, 2020. 136(9): p. 1023–1032.

8. Savani, B., et al., Management of relapse after allo-SCT for AML and the role of second transplantation. Bone marrow transplantation, 2009. 44(12): p. 769–777.

9. Romero, A., et al., Post-consolidation immunotherapy with histamine dihydrochloride and interleukin-2 in AML. Scandinavian Journal of Immunology, 2009. 70(3): p. 194–205.

10. Joshi, S.K., et al., The AML microenvironment catalyzes a stepwise evolution to gilteritinib resistance. Cancer Cell, 2021. 39(7): p. 999–1014. e8.

11. Schroeder, T., et al., Treatment of acute myeloid leukemia or myelodysplastic syndrome relapse after allogeneic stem cell transplantation with azacitidine and donor lymphocyte infusions—a retrospective multicenter analysis from the German Cooperative Transplant Study Group. Biology of Blood and Marrow Transplantation, 2015. 21(4): p. 653–660.

12. Bertsimas, D. and H. Wiberg, Machine learning in oncology: methods, applications, and challenges. JCO Clinical Cancer Informatics, 2020. 4.

13. Radak, M., H.Y. Lafta, and H. Fallahi, Machine learning and deep learning techniques for breast cancer diagnosis and classification: a comprehensive review of medical imaging studies. Journal of Cancer Research and Clinical Oncology, 2023.

14. Karami, K., et al., Survival prognostic factors in patients with acute myeloid leukemia using machine learning techniques. PloS one, 2021. 16(7): p. e0254976.

15. Eckardt, J.-N., et al., Application of machine learning in the management of acute myeloid leukemia: current practice and future prospects. Blood Advances, 2020. 4(23): p. 6077–6085.

16. Freiesleben, J., J. Keim, and M. Grutsch, Machine learning and Design of Experiments: Alternative approaches or complementary methodologies for quality improvement? Quality and Reliability Engineering International, 2020. 36(6): p. 1837–1848.

17. Holm, E.A., et al., Overview: Computer vision and machine learning for microstructural characterization and analysis. Metallurgical and Materials Transactions A, 2020. 51: p. 5985–5999.

18. Uddin, K.M.M., et al., Machine learning-based diagnosis of breast cancer utilizing feature optimization technique. Computer Methods and Programs in Biomedicine Update, 2023. 3: p. 100098.

19. Li, J., et al., A multicenter random forest model for effective prognosis prediction in collaborative clinical research network. Artificial intelligence in medicine, 2020. 103: p. 101814.

20. Kornblau, S.M., et al., Functional proteomic profiling of AML predicts response and survival. Blood, 2009. 113(1): p. 154–64.

21. Lee, C. and G.G. Lee, Information gain and divergence-based feature selection for machine learning-based text categorization. Information processing & management, 2006. 42(1): p. 155–165.

22. Jia, P., J.-h. Dai, and Y.-h. Pan, Novel algorithm for attribute reduction based on mutual-information gain ratio. Journal-Zhejiang university engineering science, 2006. 40(6): p. 1041.

23. Manek, A.S., P.D. Shenoy, and M.C. Mohan, Aspect term extraction for sentiment analysis in large movie reviews using Gini Index feature selection method and SVM classifier. World wide web, 2017. 20: p. 135–154.

24. Haryanto, A.W. and E.K. Mawardi. Influence of word normalization and chi-squared feature selection on support vector machine (svm) text classification. in 2018 International seminar on application for technology of information and communication. 2018. IEEE.

25. Karegowda, A.G., A. Manjunath, and M. Jayaram, Comparative study of attribute selection using gain ratio and correlation based feature selection. International Journal of Information Technology and Knowledge Management, 2010. 2(2): p. 271–277.

26. Urbanowicz, R.J., et al., Relief-based feature selection: Introduction and review. Journal of biomedical informatics, 2018. 85: p. 189–203.

27. Löw, F., et al., Impact of feature selection on the accuracy and spatial uncertainty of per-field crop classification using support vector machines. ISPRS journal of photogrammetry and remote sensing, 2013. 85: p. 102–119.

28. Gislason, P.O., J.A. Benediktsson, and J.R. Sveinsson. Random forest classification of multisource remote sensing and geographic data. in IGARSS 2004. 2004 IEEE International Geoscience and Remote Sensing Symposium. 2004. IEEE.

29. Luo, X., et al., Decision-tree-initialized dendritic neuron model for fast and accurate data classification. IEEE Transactions on Neural Networks and Learning Systems, 2021. 33(9): p. 4173–4183.

30. Park, H.-A., An introduction to logistic regression: from basic concepts to interpretation with particular attention to nursing domain. Journal of Korean Academy of Nursing, 2013. 43(2): p. 154–164.

31. Pattekari, S.A. and A. Parveen, Prediction system for heart disease using Naïve Bayes. International Journal of Advanced Computer and Mathematical Sciences, 2012. 3(3): p. 290–294.

32. Guelman, L., Gradient boosting trees for auto insurance loss cost modeling and prediction. Expert Systems with Applications, 2012. 39(3): p. 3659–3667.

33. Feng, J., et al., The deubiquitinating enzyme USP20 regulates the stability of the MCL1 protein. Biochemical and Biophysical Research Communications, 2022. 593: p. 122–128.

34. Al-Kaabneh, B., B. Frisch, and O.S. Aljitawi, The Potential Role of 3D In Vitro Acute Myeloid Leukemia Culture Models in Understanding Drug Resistance in Leukemia Stem Cells. Cancers, 2022. 14(21): p. 5252.

35. De Blasio, A., et al., Loss of MCL1 function sensitizes the MDA-MB-231 breast cancer cells to rh-TRAIL by increasing DR4 levels. Journal of Cellular Physiology, 2019. 234(10): p. 18432–18447.

36. Zhang, S., et al., CDK7 inhibition induces apoptosis in acute myeloid leukemia cells and exerts synergistic antileukemic effects with azacitidine in vitro and in vivo. Leukemia & Lymphoma, 2023: p. 1–12.

37. Wu, J., et al., Study of clinical outcome and prognosis in pediatric core binding factor-acute myeloid leukemia. Zhonghua xue ye xue za zhi= Zhonghua xueyexue zazhi, 2019. 40(1): p. 52–57.

38. Lee, J.H., et al., A Novel KIT INDEL Mutation in Acute Myeloid Leukemia With t (8; 21)(q22; q22); RUNX1-RUNX1T1. Annals of Laboratory Medicine, 2016. 36(4): p. 371.

39. Cairoli, R., et al., Old and new prognostic factors in acute myeloid leukemia with deranged core-binding factor beta. American Journal of Hematology, 2013. 88(7): p. 594–600.

40. Kornblau, S.M., et al., Highly phosphorylated FOXO3A is an adverse prognostic factor in acute myeloid leukemia. Clinical Cancer Research, 2010. 16(6): p. 1865–1874.

41. Hasle, H., et al., Gemtuzumab ozogamicin as postconsolidation therapy does not prevent relapse in children with AML: results from NOPHO-AML 2004. Blood, The Journal of the American Society of Hematology, 2012. 120(5): p. 978–984.

42. Patel, V.M., et al., Duvelisib treatment is associated with altered expression of apoptotic regulators that helps in sensitization of chronic lymphocytic leukemia cells to venetoclax (ABT-199). Leukemia, 2017. 31(9): p. 1872–1881.

43. Liu, X., et al., Ubenimex suppresses the ability of migration and invasion in gastric cancer cells by alleviating the activity of the CD13/NAB1/MAPK pathway. Cancer Management and Research, 2021: p. 4483–4495.

44. Sun, L., et al., Activation of tyrosine metabolism in CD13+ cancer stem cells drives relapse in hepatocellular carcinoma. Cancer Research and Treatment: Official Journal of Korean Cancer Association, 2020. 52(2): p. 604–621.

45. Zhang, W. and K. Zhang, A transcriptomic signature for prostate cancer relapse prediction identified from the differentially expressed genes between TP53 mutant and wild-type tumors. Scientific Reports, 2022. 12(1): p. 10561.

46. Biganzoli, L. and A. McCartney, Neoadjuvant nab-paclitaxel in breast cancer: who stands to benefit? J Clin Oncol, 2019. 37: p. 2226–34.

47. Joensuu, K., et al., ER, PR, HER2, Ki-67 and CK5 in early and late relapsing breast cancer— Reduced CK5 expression in metastases. Breast cancer: basic and clinical research, 2013. 7: p. BCBCR. S10701.

